# Expression Profile of Human Bitter Taste Receptors in Human Aortic and Coronary Artery Endothelial Cells

**DOI:** 10.1101/2024.10.31.621361

**Authors:** Paweł Kochany, Mikołaj Opiełka, Paulina Słonimska, Jacek Gulczynski, Tomasz Kostrzewa, Andrzej Łoś, Łukasz Znaniecki, Agnieszka Mickiewicz, Ewa Iżycka-Świeszewska, Paweł Sachadyn, Michał Woźniak, Marcin Gruchała, Magdalena Górska-Ponikowska

## Abstract

Human bitter taste receptors (TAS2Rs) are G protein-coupled receptors primarily associated with bitter taste perception, in the oral cavity. Many dietary compounds and drugs function as TAS2Rs agonists, activating their intracellular signaling pathways. Emerging evidence indicates TAS2Rs expression in extraoral tissues, including the cardiovascular system, where their functional role remains underexplored.. This study investigates TAS2Rs expression in primary human aortic and coronary artery endothelial cells. Using digital PCR (dPCR), we confirmed the mRNA expression of all 25 TAS2R subtypes in both cell lines. The expression levels were substantially higher in human coronary artery endothelial cells compared to human aortic endothelial cells. We confirmed the protein expression of TAS2R10 and TAS2R38 proteins using Western blot. These findings mark the first identification of TAS2Rs in these endothelial cells, suggesting a potential role in vascular physiology. These findings suggest a potential role for TAS2Rs in vascular physiology, prompting further research into their impact on endothelial function in cardiovascular health and pathology.

## Introduction

Humans recognize five basic tastes: sweet, sour, bitter, salty, and umami. Bitter taste receptors (TAS2Rs) were primarily discovered in type II taste receptor cells in the tongue taste buds, where they elicit the sensation of bitterness [1]. TAS2Rs are G protein-coupled receptors (GPCR) with 25 distinct, functional subtypes identified in humans [2]. Various natural and synthetic compounds, including nutrients, toxins, and drugs, evoke a bitter taste response through TAS2Rs in humans [3]. The binding of TAS2R agonists induces a conformational change in the receptor, allowing interaction with heterotrimeric G protein – gustducin (Gαgustducin/β/γ13). However, current evidence proves that TAS2Rs can also signal through noncanonical G proteins other than gustducin [4]. The disassociation of the Gαgustducin (Gαgust) activates activates phosphodiesterases (PDE), leading to a reduction in cyclic AMP (cAMP) levels. Meanwhile, the disassociation of the Gβ/γ13 subunit triggers an intracellular signaling cascade involving phospholipase C-beta 2 (PLCβ2) resulting in the generation of Inositol 1,4,5-trisphosphate (IP3). IP3 activates the inositol trisphosphate receptor isoform 3 (IP3R3) on the surface of the endoplasmic reticulum (ER), releasing stored calcium ions. This leads to activation of the plasma membrane-localized cation channels, causing depolarization of cellular membrane potential [5, 6].

Recent studies identified TAS2Rs in numerous extraoral tissues, including upper and lower airways, gastrointestinal tract, genitourinary system, nervous system, immune cells, cardiovascular system, and cancer cells [4, 7]. However, despite their broad distribution, the precise functional roles of TAS2Rs outside the oral cavity remain poorly understood. In the cardiovascular system, TAS2Rs have primarily been studied in the heart and the vascular muscle layer [8-14]. TAS2Rs have been identified in the left ventricle and sinoatrial node of the heart, where their activation by agonists leads to negative chronotropic and inotropic effects, as demonstrated in isolated murine heart models [8, 9].

Studies on different types of arterial tissues and cell lines from both animals and humans confirmed the expression of TAS2Rs in the vascular smooth muscle cells (VSMC) [10–14]. Functional experiments have demonstrated the contrasting effects of TAS2Rs agonists on arterial tone. Depending on the vascular bed, the specific TAS2Rs agonist used, and the experimental model, both vasodilation and vasoconstriction have been observed [11–14]. The impact of TAS2Rs on heart contractility and vascular tone may be significant for cardiovascular physiology, particularly for blood pressure (BP) regulation. In rats, intravenous injection of nonselective TAS2Rs agonist - denatonium benzoate, resulted in a significant and rapid decrease in BP [10]. However, this effect may not be solely attributed to TAS2Rs activation.

The vascular endothelium plays a crucial role in cardiovascular physiology, as it regulates vascular tone, influences vascular remodeling, produces cytokines, and manages hemostasis. While TAS2Rs have been identified in the vascular endothelium, their specific role in this tissue remains poorly understood. Immunofluorescence imaging has demonstrated TAS2R7 expression in the endothelium of the rat mesenteric artery and human omental artery [13]. Additionally, several subtypes of TAS2RS were also identified in human pulmonary arterial endothelial cells (HPAECs), with TAS2R10, TAS2R14, and TAS2R38 showing the highest expression levels. Furthermore activation of TAS2R10 and TAS2R38 by agonists (denatonium benzoate and phenylthiourea, respectively) attenuated lipopolysaccharide (LPS) - induced permeability and VE-cadherin internalization in HPAECs [15].

The presence and function of TAS2Rs in other vessel types of the vascular bed, particularly in the endothelium, have not been explored. The aorta and coronary arteries are two distinct types of arteries with different physiological roles: the aorta, as the main conduit for blood from the heart to the systemic circulation, and the coronary arteries, which supply oxygenated blood to the heart itself. This study aims to investigate, for the first time, the TAS2Rs mRNA and protein expression in human aortic and coronary artery endothelium cell lines - Human Aortic Endothelial Cells (HAEC) and Human Coronary Artery Endothelial Cells (HCAEC). By identifying TAS2R expression levels in these cells, we provide a foundation for future studies exploring their function in vascular physiology and cardiovascular health.

## Materials and Methods

### Cell Culture

Primary, Normal Human Aortic Endothelial Cells (HAEC, ATCC PCS-100-011) and Primary, Normal Human Coronary Artery Endothelial Cells (HCAEC, ATCC PCS-100-020) were purchased from ATCC. The cells were grown in Vascular Cell Basal Medium (ATCC PCS-100-030) supplemented with Endothelial Cell Growth Kit-BBE (ATCC PCS-100-040), containing 2% FBS, 10 mM L-glutamine, with additional supplementation of 1% penicillin/streptomycin (Sigma-Aldrich) at 37°C with 5% CO_2_. The cells used for all experiments were from passages 4–8.

### RNA Extraction and Digital PCR

Total RNA was extracted from HAEC and HCAEC (from passages 4-5) using RNeasy Mini Kit (Qiagen, Hilden, Germany) following the manufacturer’s instructions. RNA concentration and purity were assessed using a NanoDrop2000 Spectrometer (Thermo Fischer Scientific, the U.S.A) and reverse transcripted to the complementary DNA (cDNA) using TranScriba Kit (A&A Biotechnology). Gene expression was measured using specific, validated PCR TaqMan™ primers (FAM-MGB linked, Thermo Fisher Scientific, the U.S.A) (S1 Table). Digital polymerase chain reaction (dPCR) was performed using the QIAcuity Digital PCR System (Qiagen, Hilden, Germany), according to the manufacturer’s instructions. The total reaction mix (12μl) was transferred to 96-well QIAcuity Nanoplate 8.5k (Qiagen, Hilden, Germany) and then sealed. Each sample was analyzed in duplicate. Partitioning, cycling and imaging were performed in the QIAcuity One, 2plex Device (Qiagen, Hilden, Germany). The thermal cycling conditions were as follows: initial heat activation – 2 minutes, 95 °C, 40 two-step cycles – 2 seconds, 95 °C, followed by 30 seconds, 60 °C. TAS2Rs mRNA absolute values were referred to the reference gene - TATA-box binding protein (TBP). Data are presented from two independent dPCR experiments.

### Western Blot

The protein levels of TAS2R10 (PA5-102248, Thermo Fischer Scientific, the U.S.A), TAS2R38 (OST00440W-100UL, Thermo Fischer Scientific, the U.S.A) and β-actin (A3854, Sigma-Aldrich, Poland) were detected in HAEC and HCAEC cell lines using the Western blot method. The cells were seeded on 75 cm2 culture flasks and then, at confluence > 80% were scraped out and centrifuged. The pellets were suspended in RIPA buffer (R0278, Sigma-Aldrich, Poland) and a protease inhibitor cocktail – EASY pack cOmplite, EDTA-Free (Roche, 5892791001, Sigma-Aldrich, Poland). Protein concentration was determined using the Bradford reagent (Sigma-Aldrich, Poland). After that, samples containing 40 μg of protein were combined with Laemmli loading buffer (Sigma-Aldrich, Poland) and incubated at 95 °C for 5 minutes. Electrophoresis was conducted at 120 mV for 90 minutes to separate the proteins. The separated proteins were then transferred to Trans-Blot Turbo Midi 0.2 μm PVDF membranes (1704157, Bio-rad, the U.S.A) using the Trans-Blot Turbo System (2,5 A, 25 V for 7 minutes). After blocking with 5% nonfat milk in TBS-T (0.5% Tween20, 20 mM Tris-HCl, pH 7.4, and 0.5 M NaCl) for 45 minutes, the membranes were incubated with primary antibodies overnight at 4 °C. After being washed 3 times for 10 minutes each in TBS-T, the membranes were incubated with horseradish peroxidase (HRP-conjugated secondary antibodies, 1:50,000 dilution in TBS-T) for 1 h at room temperature (Anti-rabbit IgG peroxidase antibody produced in goat, A0545, Sigma Aldrich). Subsequently, the membranes were washed three times for ten minutes each in TBS-T. Chemiluminescence visualization was performed according to the manufacturer’s instructions using LuminataTM Crescendo Western HRP Substrate (Millipore Corporation, Burlington, MA).

### Data Processing and Statistical Analysis

The dPCR data were processed using QIAcuity Software Suite (Qiagen, Hilden, Germany). The number of transcripts was calculated using Poisson statistics based on the total number of positive partitions. The statistical significance of the differences between TAS2Rs relative mRNA expression in HAEC and HCAEC were carried out using one-way ANOVA followed by post-hoc Tukey tests) with XLSTAT software (Addinsoft). Graphs were processed using GraphPad Prism software (version 8, GraphPad Software, Inc., San Diego, CA, the U.S.A). Statistical significance was marked as follows: *p < 0.05, **p < 0.01.

## Results and Discussion

To the best of our knowledge, this is the first study that identifies TAS2Rs expression in human aortic and coronary artery endothelial cells. Digital PCR (dPCR) was conducted to determine the absolute expression levels of 25 human TAS2R subtypes mRNA within HAEC and HCAEC cell lines. Compared to well-established real-time quantitative PCR (RT-qPCR) using light cyclers, dPCR offers greater precision and sensitivity, particularly for low-abundant targets [16, 17]. The dPCR technique allows precise measurement of the transcript numbers in a sample, achieved by distributing cDNA molecules into thousands of nanoplate partitions. As a result, an individual partition represents a single transcript and is analyzed separately. Therefore, despite the low number of copies, we successfully obtained mRNA expression data for all 25 functional TAS2R subtypes at a stable level. The results were obtained as an absolute quantification of target gene copies. The analysis of each replicate was valid, within confidence intervals. TAS2Rs mRNA values were referred to the reference gene – TBP (Fig 1).

**Fig 1.**
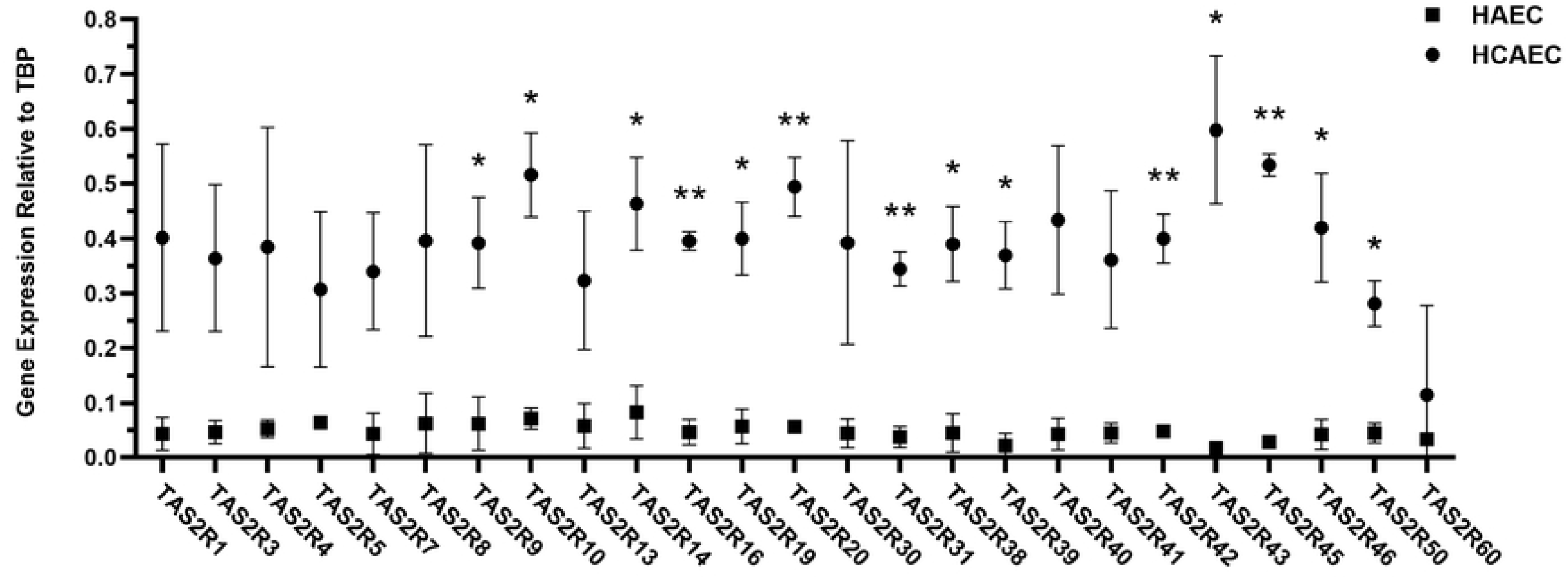
The mRNA expression of TAS2R subtypes in HAEC and HCAEC. The mRNA expression of TAS2R subtypes demonstrated as relative to TBP (which was normalized as 1), in HAEC and HCAEC. Data are presented as mean ± SD, n=2, * p < 0.05; ** p < 0,01.

The most expressed TAS2R subtypes in HAEC were TAS2R14, TAS2R10, and TAS2R5 and in HCAEC - TAS2R43, TAS2R45, and TAS2R10. Interestingly, the expression of all TAS2R subtypes was higher in HCAEC compared to HAEC (Table 1). We believe this difference may arise from the distinct physiological roles of TAS2Rs in these arteries.

**Table 1.** The mRNA expression of TAS2R subtypes in arterial endothelium.

| Bitter taste receptor<br>(TAS2Rs) | mRNA expression<br>in HAEC<br>(relative to *TBP*) ± SD | mRNA expression<br>in HCAEC<br>(relative to *TBP*) ± SD | Ratio of mRNA<br>expression in<br>HCAEC/HAEC | mRNA expression<br>in HPAECs<br>(relative to *TBP*)<br>± SD [15] |
| --- | --- | --- | --- | --- |
| *TAS2R1* | 0.04 ± 0.03 | 0.40 ± 0.17 | 9.25 | 0.49 ± 0.46 |
| *TAS2R3* | 0.05 ± 0.02 | 0.36 ± 0.13 | 7.84 | 0.49 ± 0.23 |
| *TAS2R4* | 0.05 ± 0.02 | 0.38 ± 0.22 | 7.27 | 0.82 ± 0.55 |
| *TAS2R5* | 0.06 ± 0.001 | 0.31 ± 0.14 | 4.76 | n.d. |
| *TAS2R7* | 0.04 ± 0.04 | 0.34 ± 0.11 | 7.83 | n.d. |
| *TAS2R8* | 0.06 ± 0.06 | 0.40 ± 0.18 | 6.32 | n.d. |
| *TAS2R9* | 0.06 ± 0.05 | 0.39 ± 0.08 | 6.30 * | n.d. |
| *TAS2R10* | 0.07 ± 0.02 | 0.52 ± 0.08 | 7.21 * | 2.09 ± 0.37 |
| *TAS2R13* | 0.06 ± 0.04 | 0.32 ± 0.13 | 5.56 | n.d. |
| *TAS2R14* | 0.08 ± 0.05 | 0.46 ± 0.08 | 5.59 * | 2.41 ± 0.48 |
| *TAS2R16* | 0.05 ± 0.02 | 0.40 ± 0.02 | 8.45 ** | 0.52 ± 0.29 |
| *TAS2R19* | 0.06 ± 0.03 | 0.40 ± 0.07 | 6.98 * | n.d. |
| *TAS2R20* | 0.06 ± 0.01 | 0.49 ± 0.05 | 8.77 ** | n.d. |
| *TAS2R30* | 0.04 ± 0.03 | 0.39 ± 0.19 | 8.78 | n.d. |
| *TAS2R31* | 0.03 ± 0.02 | 0.34 ± 0.03 | 9.14 ** | n.d. |
| *TAS2R38* | 0.05 ± 0.04 | 0.39 ± 0.07 | 8.63 * | 2.43 ± 0.29 |
| *TAS2R39* | 0.02 ± 0.02 | 0.37 ± 0.06 | 16.92 * | n.d. |
| *TAS2R40* | 0.04 ± 0.03 | 0.43 ± 0.14 | 10.09 | 0.35 ± 0.49 |
| *TAS2R41* | 0.05 ± 0.02 | 0.36 ± 0.13 | 8.02 | n.d. |
| *TAS2R42* | 0.05 ± 0.01 | 0.40 ± 0.04 | 8.33 ** | n.d. |
| *TAS2R43* | 0.01 ± 0.01 | 0.60 ± 0.14 | 34.24 * | 0.12 ± 0.28 |
| *TAS2R45* | 0.03 ± 0.01 | 0.53 ± 0.02 | 18.86 ** | n.d. |
| *TAS2R46* | 0.04 ± 0.03 | 0.42 ± 0.10 | 9.92 * | n.d. |
| *TAS2R50* | 0.05 ± 0.02 | 0.28 ± 0.04 | 6.22 * | n.d. |
| *TAS2R60* | 0.03 ± 0.01 | 0.11 ± 0.16 | 3.42 | n.d. |

Expression demonstrated as relative to TBP (which was normalized as 1) in HAEC and
HCAEC. Data are presented as mean, ± SD, n=2. Additionally, the table presents the
fold difference of TAS2R mRNA expression between HAEC and HCAEC. For
comparison, the mRNA expression of TAS2Rs relative to TBP in HPAECs is also
included from the external literature [15].
\* $p < 0,05$
\*\* $p < 0,01$
n.d. (No data) – mRNA expression undetected [15].

Kertesz et al [15] showed the expression of TAS2Rs in HPAECs. Using RT-qPCR they identified mRNA of 9 TAS2R subtypes, while the remaining subtypes were undetected (Table 1). TAS2Rs with available data in HPAECs, such as TAS2R10, TAS2R14, and TAS2R38, show relatively higher expression in HPAECs compared to both HAEC and HCAEC. For instance, TAS2R38 expression is notably higher in HPAECs (2.43 ± 0.29) than in HAEC (0.05 ± 0.04) and HCAEC (0.39 ± 0.07), indicating a potential distinct functional role for TAS2R38 in pulmonary circulation. Additionally, TAS2R45 exhibits a significant increase in expression in HCAEC (18.86-fold) compared to HAEC, while no expression was detected in HPAECs, suggesting a more specialized role in coronary circulation. Given the higher expression of TAS2Rs in coronary endothelial cells, particularly TAS2R43 and TAS2R45, these receptors may influence blood flow regulation in response to bitter compounds, which could be relevant in cardiovascular diseases. However, it is important to note that protein levels do not always directly correlate with transcript levels. This variance can result from posttranscriptional regulation, differences in translation efficiency, or protein degradation. Functional experiments in the future are needed to fully understand the physiological role of TAS2Rs. Our use of dPCR enabled the identification of all TAS2R subtypes in coronary and aortic endothelium opening the opportunity to study their functional role and signaling mechanism in the future.

To investigate the presence of TAS2R proteins in HAEC and HCAEC cell lysates, Western blotting was performed. Two TAS2R subtypes were selected for analysis: TAS2R10 and TAS2R38. TAS2R10 demonstrated significant mRNA expression in both cell lines and has been previously described in cardiovascular contexts while TAS2R38 is recognized for its role in inducing nitric oxide production [4,18-22]. The results support the dPCR findings, confirming TAS2Rs expression both on mRNA and protein levels (Fig 2).

**Fig 2.**
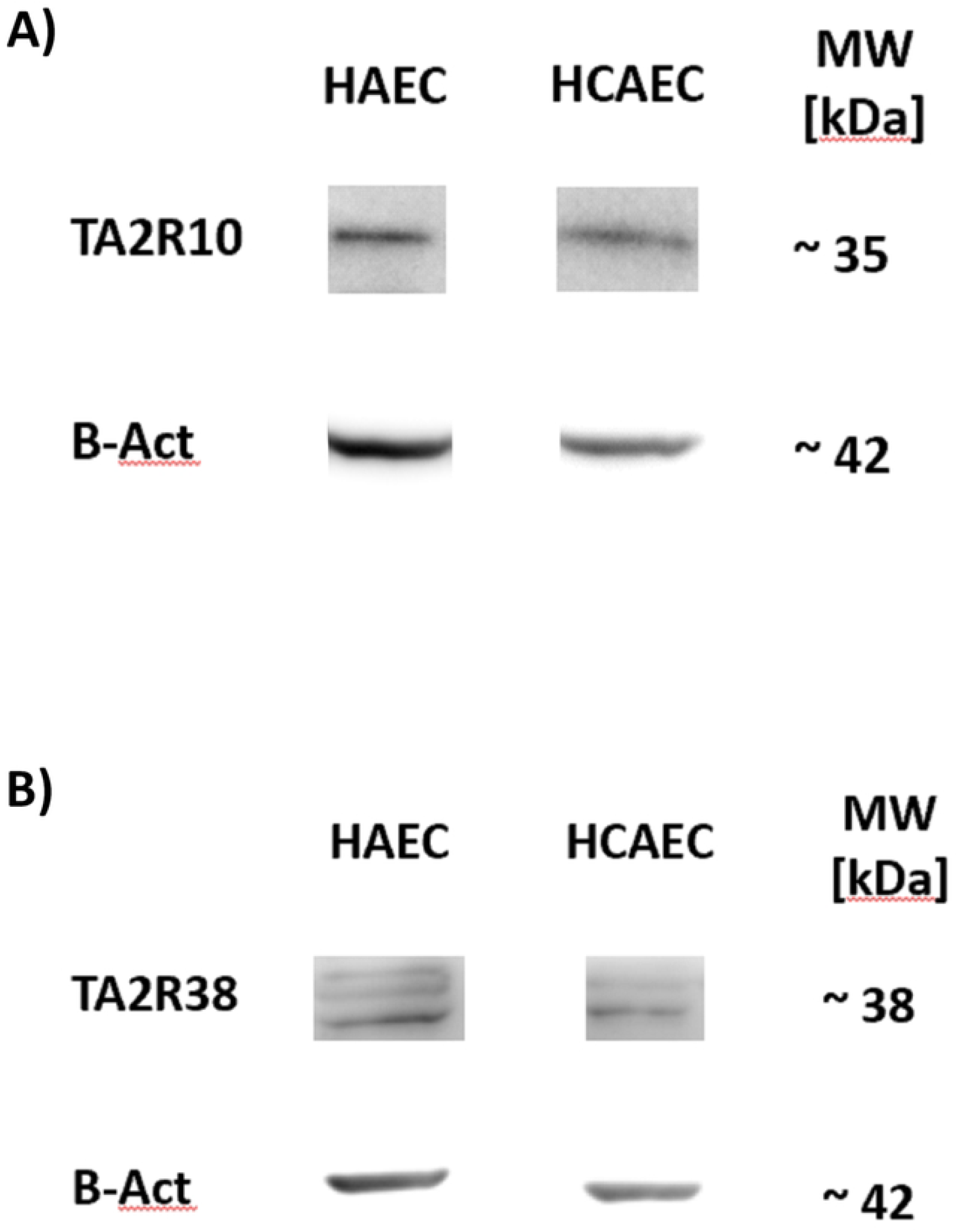
The protein expression of TAS2R subtypes in HAEC and HCAEC. The protein expression of (A) TAS2R10 and (B) TAS2R38 assessed by Western blot analysis. Representative immunoblots from one membrane are shown.

The aversive sensation of bitterness is thought to serve as a protective mechanism to prevent ingesting potential toxins. Nonetheless, humans frequently consume bitter substances, including medications and dietary compounds. Apart from the taste sensation, those substances can cause systemic effects and influence overall health. The cardiovascular effects of various TAS2Rs agonists have been documented previously, although not in the context of selective TAS2Rs activation. Examples include caffeine, epigallocatechin gallate, resveratrol, and pharmaceuticals used in clinical practice, like azithromycin, dextromethorphan, chloroquine, and diphenhydramine [4]. Endothelium-independent relaxation of precontracted human pulmonary arteries has been observed in response to TAS2R agonists [12]. In contrast, denatonium benzoate increased the tone of endothelium-denuded aorta rings in a rat model, linked to the activation of specific TAS2Rs [14]. Exploring TAS2Rs’ molecular mechanisms in coronary and aortic circulation could enhance our understanding of how bitter compounds in food and medications affect endothelial function. This insight may also help mitigate adverse drug effects.

Previous studies on TAS2Rs role in extraoral locations suggested several functional mechanisms of action. One of the most important functions is their ability to induce nitric oxide (NO) synthesis. Activation of TAS2Rs in airway epithelial cells and macrophages by various agonists, including bacterial quorum-sensing molecules of Pseudomonas aeruginosa, induces intracellular calcium [Ca2+] signaling, resulting in nitric oxide synthase (NOS) induction [19–22]. NO is one of the most important signaling molecules in the cardiovascular system. There are three isoforms of NOS: neuronal NOS (nNOS), inducible NOS (iNOS), and endothelial NOS (eNOS) [23]. All three isoforms are found in the endothelium, with eNOS being the predominant form. However, only nNOS and eNOS are Ca2+ dependent. Endothelium-derived NO is essential for maintaining vascular homeostasis and protecting the endothelium. It achieves this primarily by regulating blood flow and lowering BP through vasodilation. Moreover, NO plays a role in vascular protection against thrombosis and atherosclerosis by inhibiting platelet aggregation and leukocyte adhesion to the vascular wall [24]. NO also contributes to a complex role in maintaining redox balance. At low concentrations, NO functions as an antioxidant, helping to reduce the production of reactive oxygen species (ROS). However, when NO levels are elevated, it can react with superoxide to form peroxynitrite, which contributes to increased oxidative stress within cells, potentially leading to harmful pathological effects [25]. Moreover, NO plays a key role in promoting angiogenesis by stimulating endothelial cell proliferation, migration, and survival at low concentrations. However, at higher concentrations, NO can inhibit angiogenesis by inducing cytostasis and cell death [26]. However, dysfunctional NO synthesis has been associated with various cardiovascular diseases constituting a leading health problem, such as hypertension, atherosclerosis, aneurysm development, myocardial remodeling, and heart failure [27]. In Fig 3, we illustrate a hypothetical pathway for TAS2Rs signaling leading to NO synthesis in endothelial cells, based on our current results and previous studies of TAS2Rs in extraoral tissues. In addition, we present a summary of the cardiovascular effects of endothelium-derived NO [4,18-22, 24-26].

**Fig 3.**
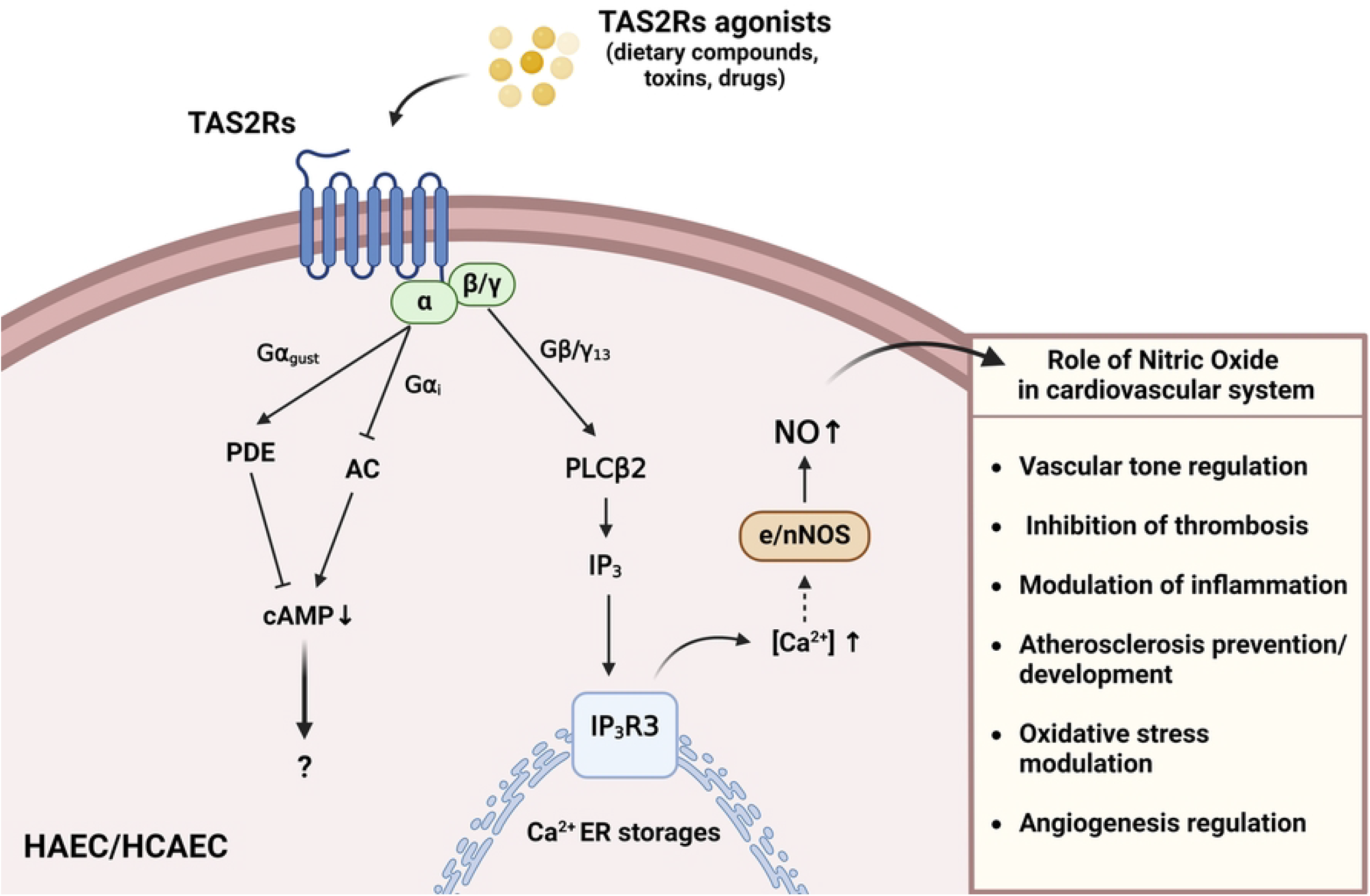
Potential TAS2Rs intracellular signaling in endothelial cells, leading to nitric oxide synthesis and impact of nitric oxide on the cardiovascular system.

As mentioned before Kertesz et al [13] showed the TAS2Rs protective function against endothelial barrier permeability induced by LPS and VE-cadherin internalization in HPAECs. LPS-induced barrier leak was associated with the reduction in cAMP and Rac1 expression in endothelial cells. TAS2R agonists (denatonium and phenylthiourea), attenuated the LPS-induced fall in cAMP levels. Notably, prior studies of TAS2R signaling, including canonical taste transduction, have shown a reduction in cAMP (Fig 3). This suggests that TAS2Rs may function through diverse, complex mechanisms, warranting further investigation.

## Conclusions

Identifying TAS2R expression in HAEC and HCAEC opens a discussion about TAS2R functions in arterial vascular endothelium. We hypothesize that activation of TAS2Rs in endothelial cells may lead to NO synthesis, impacting cardiovascular health. This topic requires further investigation to establish the physiological and pathophysiological role of TAS2RS in the endothelium.

While our study provides the first insights into TAS2R expression in human aortic and coronary artery endothelial cells, certain limitations must be acknowledged. The use of only two cell lines limits the generalizability of our findings, and further studies are needed using additional cell lines, tissue samples and in vivo studies to strengthen the conclusions. Additionally, the exact molecular mechanisms through which TAS2Rs influence endothelial function remain unclear and require future research, particularly in terms of signaling pathways such as calcium signaling and NO synthesis. Expanding on this research could pave the way for novel therapeutic options and explain how daily dietary substances and drugs impact cardiovascular health. Despite these limitations, our study lays the foundation for future exploration of TAS2Rs in vascular endothelium physiology.

## Supporting information

**S1 Table. TaqMan™ Gene Expression Assay (FAM) Primers ID used to detect human TAS2Rs in HAEC and HCAEC**.

